# Stimulus-induced gamma sources reduce in power but not in spatial extent with healthy aging in human EEG

**DOI:** 10.1101/2024.01.09.574816

**Authors:** Ankan Biswas, Wupadrasta Santosh Kumar, Kanishka Sharma, Supratim Ray

**Affiliations:** Centre for Neuroscience, Indian Institute of Science, Bengaluru, India, 560012; IISc Mathematics Initiative, Indian Institute of Science, Bengaluru, India, 560012, Facsimile

**Keywords:** EEG, Gamma oscillations, healthy aging, cluster shrinkage

## Abstract

Aging alters brain structure and function, and studying such changes may help understand the neural basis underlying aging and devise interventions to detect deviations from healthy progression. Electroencephalogram (EEG) offers an effective way to study healthy aging owing to its high temporal resolution and affordability. Recent studies have shown that narrow- band stimulus-induced gamma oscillations (20-70 Hz) in EEG, induced with cartesian gratings in a fixation task paradigm, weaken with healthy aging and onset of Alzheimer’s Disease (AD) while remaining highly reproducible for a given subject, and thus hold promise as potential biomarkers. However, functional connectivity (FC) sometimes changes in a different way compared to sensor power with aging. This difference could be potentially addressed by studying how underlying gamma sources change with aging, since either a reduction in source power or a shrinkage of the sources (or both) could reduce the power in the sensors but may have different effects on other measures such as FC. We therefore reconstructed EEG gamma sources through a linear inverse method called exact Low-resolution Tomography Analysis (eLORETA) on a large (N=217) cohort of healthy elderly subjects (>50 years). We further characterized gamma distribution in cortical space as an exponential fall-off from a seed voxel with maximal gamma source power to delineate a reduction in magnitude versus shrinkage. We found significant reduction in magnitude but not shrinkage with healthy aging. Overall, our results shed light on changes in EEG gamma source distribution with healthy aging, which could provide clues about underlying neural mechanisms.

## Introduction

Aging is a biological process that impacts brain function at multiple levels, such as cognitive, behavioural (Riddle, 2007) and physiological (Anders & Kristine, 2010). Brain aging manifests in the loss of dendritic spines, white matter density, neurotransmitter levels and the neuronal population (Hara *et al*., 2012). The study of healthy aging therefore would lead to better understanding of the changes in the brain at various levels, which could help in detecting deviations from healthy aging due to mental disorders such as Alzheimer’s Disease (AD). Aging has been studied non-invasively in various modalities, for instance in fMRI (Dennis & Thompson, 2014), EEG (Scally *et al*., 2018), and MEG (Mandal *et al*., 2018). EEG holds great promise as a diagnostic tool because it is inexpensive and provides a direct measure of postsynaptic activity and synchronization across large scale functional networks (Lopes da Silva, 2013). With healthy aging, various measures in EEG like P300 (Porcaro *et al*., 2019), N2pc (Pagano *et al*., 2015), alpha power (Borghini *et al*., 2018; Knyazeva, 2021), other resting state oscillations (Rossini *et al*., 2007) and aperiodic activity (Voytek *et al*., 2015) have been shown to alter. Recently, stimulus-induced narrow-band gamma oscillations, which are produced by presenting visual stimuli such as gratings and are thought to reflect excitatory- inhibitory interactions in the brain (Murty *et al*., 2018), have also been shown to weaken with aging (Murty *et al*., 2020). These stimulus-induced gamma oscillations are highly reproducible across recordings separated by one year or more (Kumar *et al*., 2022) and also weaken with onset of AD (Murty *et al*., 2021), making them a potentially useful biomarker.

Apart from power measured using EEG sensors, various ensemble features like functional connectivity (FC) (Kumar & Ray, 2023) and related compensatory mechanisms (Pathak *et al*., 2022) may also change with healthy aging, albeit in a different way compared to power (Murty *et al*., 2020). For example, FC decreases with aging in alpha and stimulus-induced gamma oscillations, even though alpha power does not and even when variation in gamma power is accounted for using regression (Kumar & Ray, 2023). Similarly, FC at Individual Alpha Peak Frequency (IAPF) may stay invariant with aging due to a compensatory mechanism where enhanced inter-areal coupling cancels the effect of increased axonal delays (Pathak et al., 2022). One way to further understand aging related changes in the brain is through source localization. In particular, power in sensor space can reduce if the underlying sources weaken, shrink, or a combination of the two (Figure 1), even though these may have different effects on other measures such as FC (see Discussion for more details).

**Figure 1.**
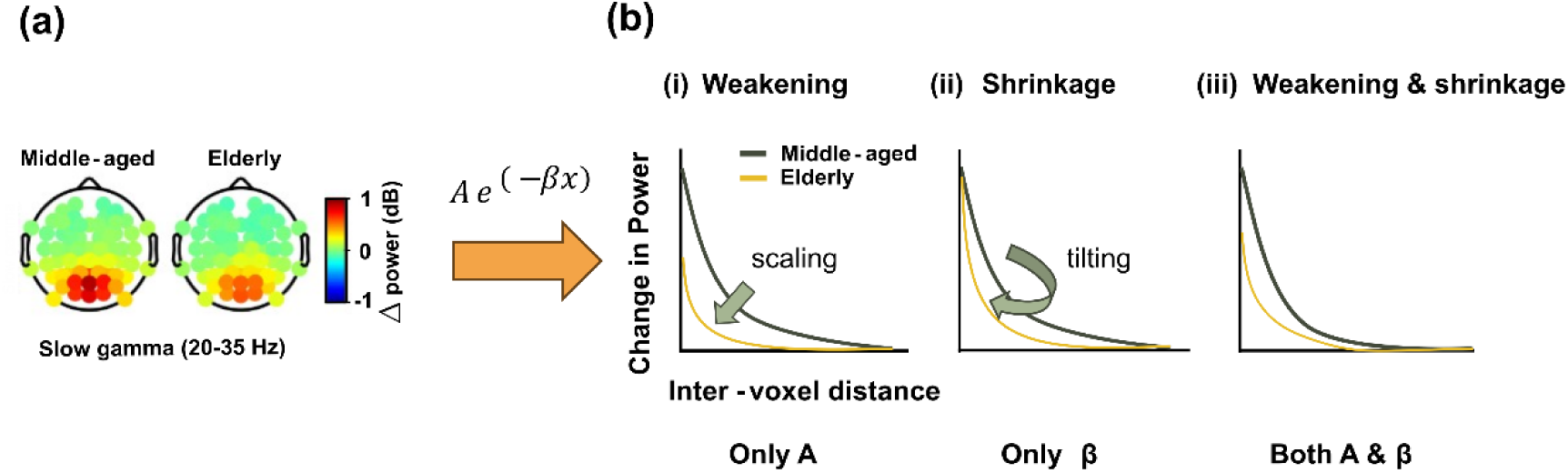
The hypothesis of changes in gamma sources based on scalp power. **(a)** The change in power scalp plot depicting the power fall-off across the occipital electrodes for middle-aged and elderly subjects as reported in (Murty *et al*., 2020) for the slow gamma range (20-35 Hz) as an illustration. Slow gamma power reduces in elderly subjects compared to the middle-aged. **(b)** Hypothetical plots describing the three possibilities for the source distribution, parametrized as exponential decay function (A*exp(-βx)), with parameters A and β. **(i)** When the source configuration remains similar, with an overall reduction in power magnitude, the profiles can be explained by the scaling factor (A), which we refer here as weakening. **(ii)** Source extent shrinkage without considerable change in seed/reference power magnitude, could be explained by the decay factor (β), which we refer here as shrinkage. **(iii)** Scenario wherein both source magnitude and the source extent changes, both the parameters A and β would be needed for explanation (combined weakening and shrinkage).

**Figure 2.**
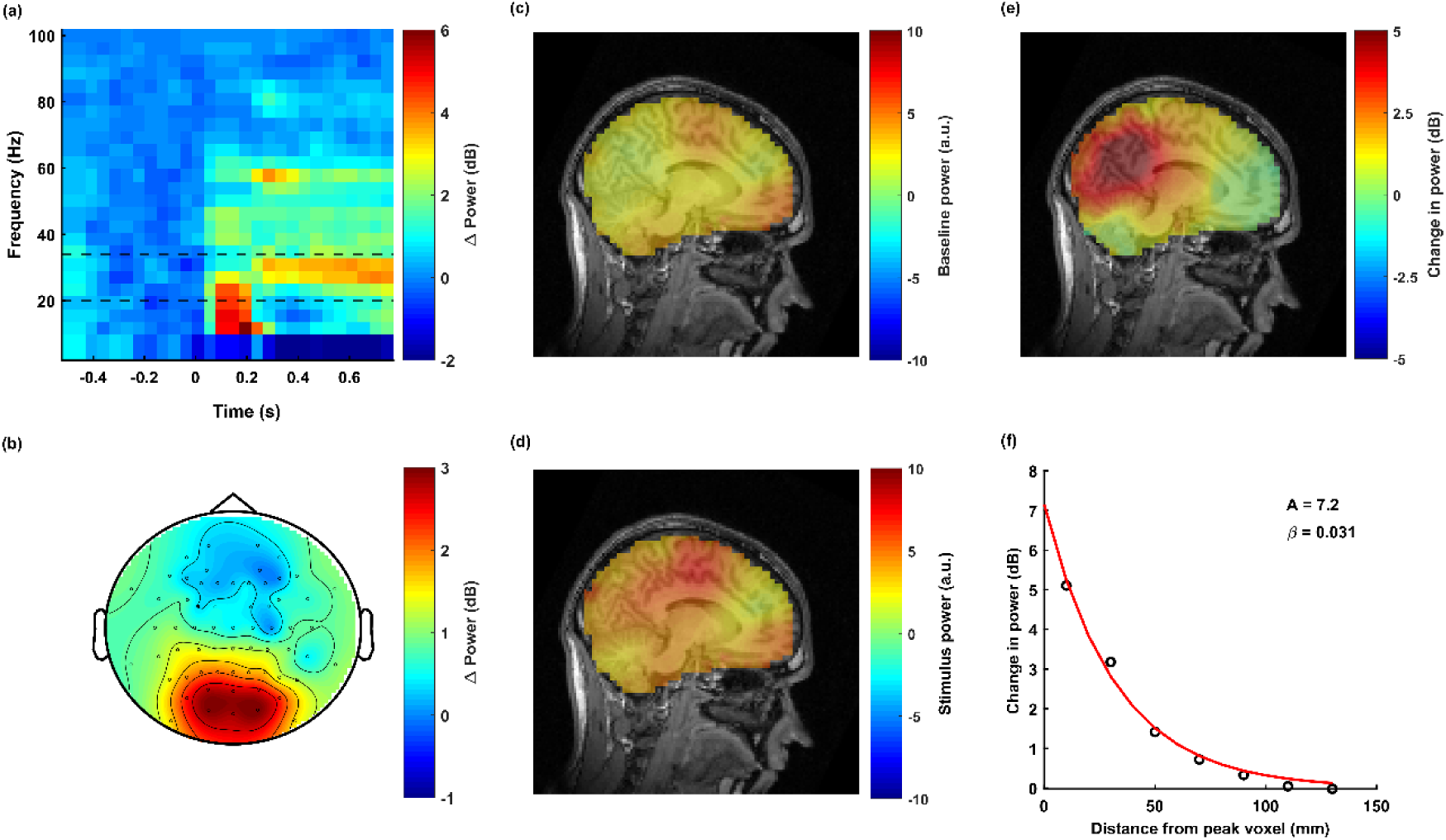
Source localization using the eLORETA method in an example subject. (a) Time-frequency change in power spectrogram averaged over occipital electrodes. The slow gamma range (20-35 Hz) is indicated by dotted black lines. (b) Corresponding topoplots for the slow gamma frequency range (c) Source distributions of baseline power (–0.5 to 0 s) for slow gamma frequency range using eLORETA. (d) Source distributions of the stimulus period (0.25 to 0.75 s). (e) Change in source power during the stimulus period compared to baseline. (f) The mean change in source power with inter-voxel distance from the peak voxel (having the highest power) is plotted using a black circle. The data are fitted with an exponential decay function (shown as a red line along with the A and β parameters).

To compare these alternatives, we considered the same EEG dataset reported in previous studies (Murty *et al*., 2020; Murty & Ray, 2022; Kumar & Ray, 2023) and performed brain source localization of visual gratings-induced gamma, along with alpha, using eLORETA (exact Low-resolution Tomography Analysis) source reconstruction method. Gamma (and alpha) sources were strongest in the occipital areas, which allowed us to model the source power as an exponential decay function (A*exp(-*β*x)) from the voxel that had maximal gamma power. Here, weakening of source power with age without any change in the distribution would result in only a reduction in A, while only a shrinkage in source distribution without any change in magnitude would increase *β*. Reduction in both magnitude and active area would change both *A* and *β*. We therefore studied changes in these parameters with aging using linear regression.

## Materials and Methods

### Human Subjects

The EEG dataset was collected from 257 human subjects (females: 106) aged 50 to 88 years as part of the Tata Longitudinal study of aging (TLSA), of which usable data was obtained from 244 subjects (227 healthy, 12 MCI and 5 AD; see (Murty *et al*., 2021) for detailed selection criteria). Subjects were recruited from urban Bengaluru communities and were evaluated by trained psychiatrists, psychologists and neurologists affiliated with National Institute of Mental Health and Neuroscience (NIMHANS) and M.S. Ramaiah Hospital, Bengaluru. Cognitive performance was evaluated using ACE (Addenbrooke’s Cognitive Examination), CDR (Clinical Dementia Rating) and HMSE (Hindi Mental State Examination) tests (see Murty et al., 2021 for more details). Since this was a community-based study, wherein subjects were recruited based on advertisements, we did not have any knowledge of their cognitive status prior to the experiment. Since the number of MCI and AD subjects were relatively few, we considered only the healthy population (N=227). We further discarded 9 healthy subjects because data was collected using 32 channels. In addition, one of the healthy subjects was discarded that had relatively fewer trials (<100) across all the protocols. All results are therefore based on the remaining 217 healthy subjects. All subjects took part against monetary compensation and provided signed informed consent. All the procedures were approved by The Institute Human Ethics Committees of Indian Institute of Science, NIMHANS, Bengaluru and M.S. Ramaiah Hospital, Bengaluru.

### Experimental Setting and Behavioural Task

Experimental setup and details of recordings have been explained in detail previously (Murty *et al*., 2020, 2021) and are summarized here. EEG was recorded from 64-channel active electrodes (actiCap) using BrainAmp DC EEG acquisition system (Brain Products GMbH) and were placed according to the international 10-10 system. Raw signals were filtered online between 0.016Hz (first-order filter) and 1kHz (fifth order Butterworth filter), sampled at 2.5kHz, and digitized at 16-bit resolution (0.1µV/bit). It was subsequently decimated to 250Hz. Average impedance of the final set of electrodes was (mean±SEM) 7.82 ± 0.02kΩ. EEG signals recorded were in reference to electrode FCz during acquisition (termed here as a “unipolar” reference scheme since all electrodes are referenced to a single electrode).

All subjects sat in a dark room facing a gamma-corrected LCD monitor (dimensions: 20.92 x 11.77 inches; resolution: 1280 x 720 pixels; refresh rate: 100 Hz; BenQ XL2411; maximum luminance of 120 cd/m^2^) with their head supported by a chin rest. It was placed ∼58 cm from the subjects and subtended 52° x 30° of visual angle for full screen gratings. Subjects performed a visual fixation task, wherein 2-3 full screen grating stimuli were presented for 800ms with an inter-stimulus interval of 700ms after a brief fixation of 1000ms in each trial using a customized software written in Objective C with OpenGL libraries running on MAC OS. The stimuli were full contrast sinusoidal luminance achromatic gratings with one of the three spatial frequencies (1, 2, and 4 cycles per degree (cpd)) and four orientations (0°, 45°, 90°, and 135°). Details of this experimental paradigm, including images of these stimuli, can be found in (Murty *et al*., 2021; Murty & Ray, 2022).

### Artifact rejection

We implemented a fully automated artifact rejection framework (for details, see (Murty *et al*., 2020, 2021; Murty & Ray, 2022), in which outliers were detected as repeats with deviation from the mean signal in either time or frequency domains by more than 6 standard deviations, and subsequently electrodes with too many (30%) outliers were discarded. Subsequently, stimulus repeats that were deemed bad in the visual electrodes (P3, P1, PO3, P2, P4, PO4, POz, O1, Oz, and O2) or in more than 10% of the other electrodes were discarded. This gave rise to a set of good electrodes and common bad repeats for each subject. We also rejected electrodes with high impedance (>25kΩ). Finally, we computed slopes for the baseline power spectrum between 56-84Hz range for each unipolar electrode and rejected electrodes whose slopes were less than 0.

Eye position was monitored using EyeLink 1000 (SR Research Ltd), sampled at 500Hz. Any eye-blinks or change in eye position outside a 5° fixation window during -0.5s to 0.75s from stimulus onset were identified as fixation breaks, which were removed offline. We have previously shown that eye-movements and microsaccades (Hassler *et al*., 2011) have a negligible effect on gamma power in this dataset (Murty *et al*., 2020). The number of repeats available for analysis were 151.1±3.7 for middle aged and 140.7±3.3 for the elderly group.

### EEG Data Analysis

Analyses were performed using the 10 electrodes in occipital lobe, namely, P3, P1, PO3, P2, P4, PO4, POz, O1, Oz, and O2. We considered the unipolar reference scheme which suitable for the source localization procedure. These electrodes were considered because strong gamma oscillations were observed in these electrodes, as done previously in our previous studies (Murty *et al*., 2020, 2021; Kumar *et al*., 2022). All the data analyses were done using custom codes written in MATLAB (MathWorks. Inc; RRID:SCR_001622). Change in power between stimulus period (ST: 250-750 ms after stimulus onset) and baseline period (BL: -500 to 0 ms, where 0 indicates stimulus onset) for a frequency band was computed using the following formula:

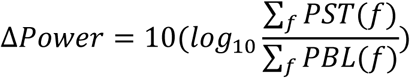

where PST is the stimulus power spectra and PBL is the baseline power spectra, both averaged within relevant frequency bins (*f*), across all the analyzable repeats. The time intervals for stimulus and baseline were chosen to avoid the transient fluctuations which have power over a broad frequency range. Power was computed in three frequency bands: alpha (8‒12 Hz), slow gamma (20‒35 Hz), and fast gamma (36‒66 Hz). The gamma frequency band limits were set based on the localization of two gamma peaks in the change in power spectral density (PSD) profile and the same limits were used in our previous studies (Murty *et al*., 2018, 2020, 2021). Average occipital power was estimated by taking mean across all the ten electrodes listed above. Scalp maps were generated using the *topoplot* function of EEGLAB toolbox ((Delorme & Makeig, 2004), RRID:SCR_007292).

### Source localization

We localized the sources after cleaning the data of any artifacts as described above. Electrodes that were labelled bad as per the criteria mentioned in the Artifact Rejection sub-section (flat PSD, noisy and high impedance electrodes) were first reconstituted for each subject by spherical spline interpolation using *pop_interp* function of EEGLAB toolbox. Intracortical distribution of electric activity was estimated from the surface EEG data using eLORETA (Pascual-Marqui, 2007, 2009; Pascual-Marqui *et al*., 2011), which is a mathematical solution of a regularized, weighted minimum-norm with three-dimensional (3D) distribution in brain volume. Even though the reconstruction has low resolution, the method promises correct localization, based on experimental data through a virtual head and a real human EEG recorded under diverse stimulus conditions (Pascual-Marqui *et al*., 2011). Also, eLORETA is very effective in mitigating false positives (Halder *et al*., 2019). The 3D solution space of the reconstructed sources had voxels with 5 mm separation. By restricting the voxels to cortical gray matter and hippocampus, we obtained 6239 voxels in a realistic head model based on The Montreal Neurologic Institute average MRI brain (MNI152) template (Mazziotta *et al*., 2001).

Artifact-free scalp interpolated EEG data was segmented into baseline (BL) and stimulus (ST) periods. Data was segmented and saved using custom MATLAB codes to generate files suitable for eLORETA software. The eLORETA software, which has been widely used to investigate cortical electrical activities, is freely downloadable from https://www.uzh.ch/keyinst/loreta and has been validated through many studies of resting state, altered state of consciousness in neuropsychiatric diseases and during various sensory stimulations (Pazo-Álvarez *et al*., 2004; Gianotti *et al*., 2008; Lehmann *et al*., 2012; Babiloni *et al*., 2013). Using the utility window, after applying average reference, we generated cross spectrum (.crss) files from spectral power matrix for the three frequency ranges (alpha: 8‒12 Hz, slow gamma: 20‒35 Hz, fast gamma: 36‒66 Hz), which were then transformed into the cortical electrical current density data as (voxel position × time × frequency ranges) by applying eLORETA source reconstruction. Finally, the cortical electrical activity was obtained by squaring the cortical electrical current density and averaging over the entire EEG segment (Pascual-Marqui, 2007; Aoki *et al*., 2023) as .slor files for each trial for the baseline and stimulus periods for individual subjects. In line with previous studies (Jatoi *et al*., 2014), we used the term source power in the present study to refer to the squared current source density. The difference in squared current source density (ΔCSD) was obtained by log subtraction of baseline from stimulus data for each voxel × trial × subject. The .slor files for each subject were converted into text files and imported to MATLAB using custom codes to generate inter-voxel separation matrix, based on the seed voxel with maximal power change (averaged across all subjects; hence the same seed voxel was used for all subjects) for each of the three frequencies and for each hemisphere. For alpha, seed voxel was the one with maximum decrease in power change, while for slow-gamma (SG) and fast-gamma (FG), voxel with maximal increase in power change was considered across all the subjects including both the age groups. We used the same seed voxel for all subjects since the individual subject seed voxel varied across the cortex and for subjects with weak gamma, the chosen voxel was not even over the occipital area, hence giving uninterpretable results. Since the seed voxel fixed, the differences across age could not simply be due to variability across subjects and differences in the choice of the seed voxel. Inter-voxel separation across the seed voxel and all other voxels was calculated using the formula.

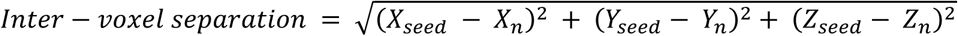

where (xseed, yseed, zseed) are the position coordinates of the seed voxel and (xn, yn, zn) are the position coordinates for other voxels in brain volume consisting of 6239 voxels.

### Simulation of the spatial extent of the source distribution

We also performed simulations to test how artificially shrinking the spatial extent of the source distribution affects data in sensor space and subsequent source localization of this sensor data (Figure 3). This required a forward model to generate the sensor data from sources, which was not directly accessible in the eLORETA software described above. We therefore, used fieldtrip toolbox (Oostenveld *et al*., 2011) in the MATLAB (MathWorks Inc., RRID: SCR_001622) to perform the simulations to estimate sources using the eLORETA technique, following a similar strategy as previous studies (Halder *et al*., 2019; Kaboodvand *et al*., 2024).

**Figure 3.**
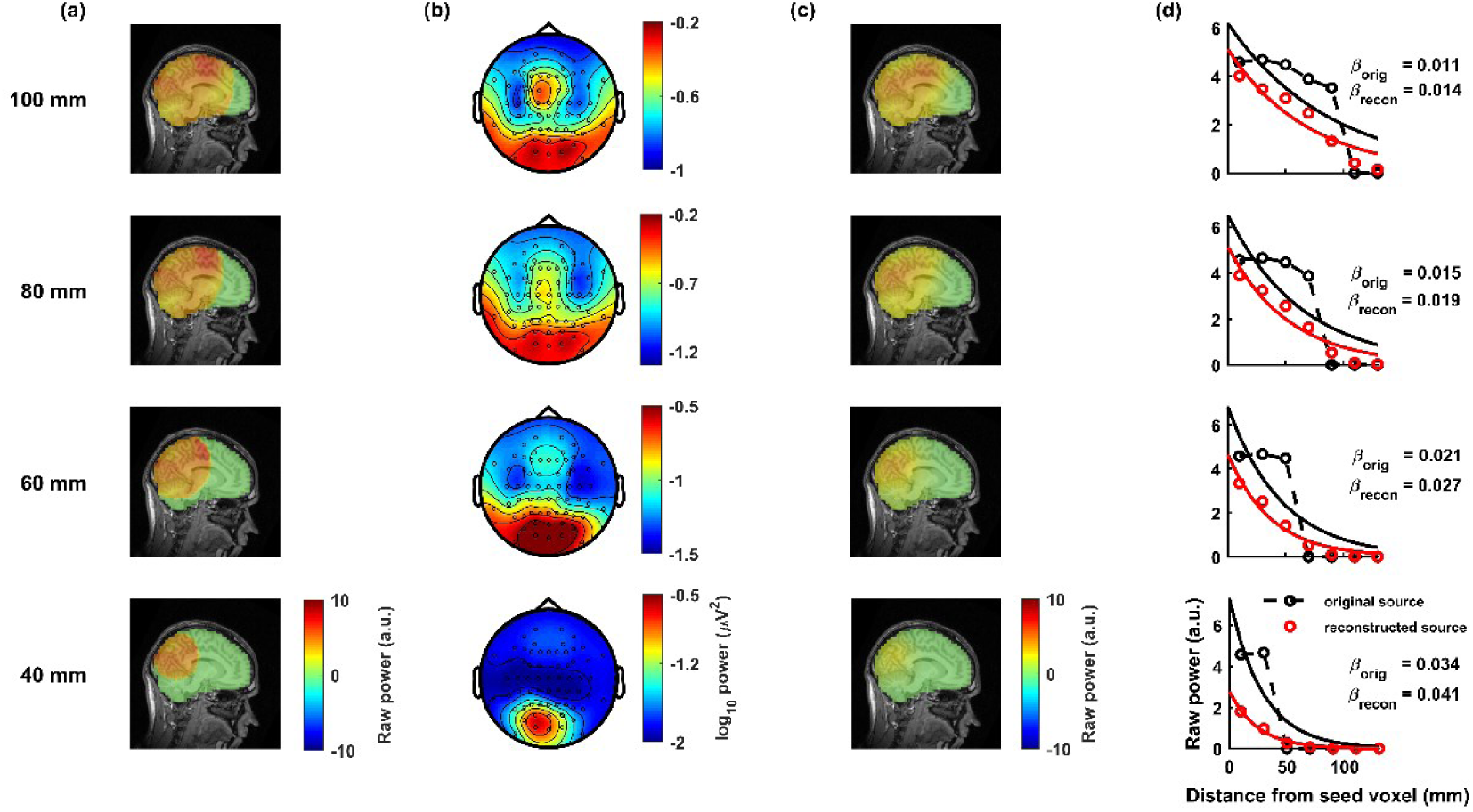
Validation of the eLORETA source localization method using simulations with varying spatial extents of source distribution (100–40 mm) in an example subject for the stimulus period. **(a)** Source maps for four different spatial distributions (100 mm, 80 mm, 60 mm, and 40 mm) constructed from the time-domain signal, shown in a sagittal cross-section of the brain. **(b)** Topoplot showing power across electrodes, computed from the time-domain signal reconstructed from the corresponding source distributions in panel (a). **(c)** Source distribution maps reconstructed from the time-domain sensor data. **(d)** The mean change in source power with inter-voxel distance from the peak voxel (having the highest power) is plotted using black circles for the original source powers (panel a) and red circles for the reconstructed source maps (panel c). The data is fitted with an exponential decay function, shown as black and red continuous lines, with the corresponding **A** and **β** values displayed in the inset.

For the simulation, we began with EEG time-series data from an example subject. The data were pre-processed using the following steps in FieldTrip: bad electrodes (see the **Artifact rejection)** were interpolated via a spline interpolation algorithm (*ft_channelrepair*), followed by average referencing (*ft_preprocessing*). The trials were then segmented into baseline and stimulus epochs using *ft_redefinetrial*. The cleaned clean time-domain data were then transformed into the frequency domain using *ft_freqanalysis*, which applied a multitaper frequency transformation to compute spectral representations (e.g., cross-spectral density matrices). Next, we constructed a head model using a standard MRI dataset in FieldTrip. This anatomical dataset was segmented and used to create a boundary element method (BEM) head model using *ft_prepare_headmodel*. To ensure accurate alignment between electrodes and the anatomical structure, we utilized *ft_electroderealign* for sensor co-registration. A lead field matrix was then computed with *ft_prepare_leadfield* for a 5-mm resolution grid-based source model.

Source activity was estimated using *ft_sourceanalysis*, with the inverse method set to ’eloreta’. This approach incorporated the cross-spectral density matrix and lead field as inputs. The eLORETA algorithm applied a spatially weighted minimum norm estimation with depth regularization to reconstruct neural sources while minimizing localization bias. Finally, the resulting source power distribution was visualized using *ft_sourceplot* to map oscillatory activity across the brain.

### Parametrizing source distribution with exponential decay function

The source map in 3D was converted to 1D scatter over all the voxels, with power against the inter-voxel separation from the voxel with maximum power. The data was binned across equally spaced inter-voxel separation. For each individual subject, exponential decay function was fit over the means in each of the bins. The exponential decay function is defined as follows.

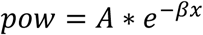

The parameters *A*, *β* were termed as the amplitude and the decay parameter, respectively. The curve fitting was done using ‘*fminsearch*’ function in MATLAB. The fall-off profiles were computed for both the hemispheres individually as depicted in Figure 4c, because there are more structural connectivity paths within each hemisphere, and that the spread profile makes sense only within a hemisphere and not crossing over the longitudinal fissure.

**Figure 4.**
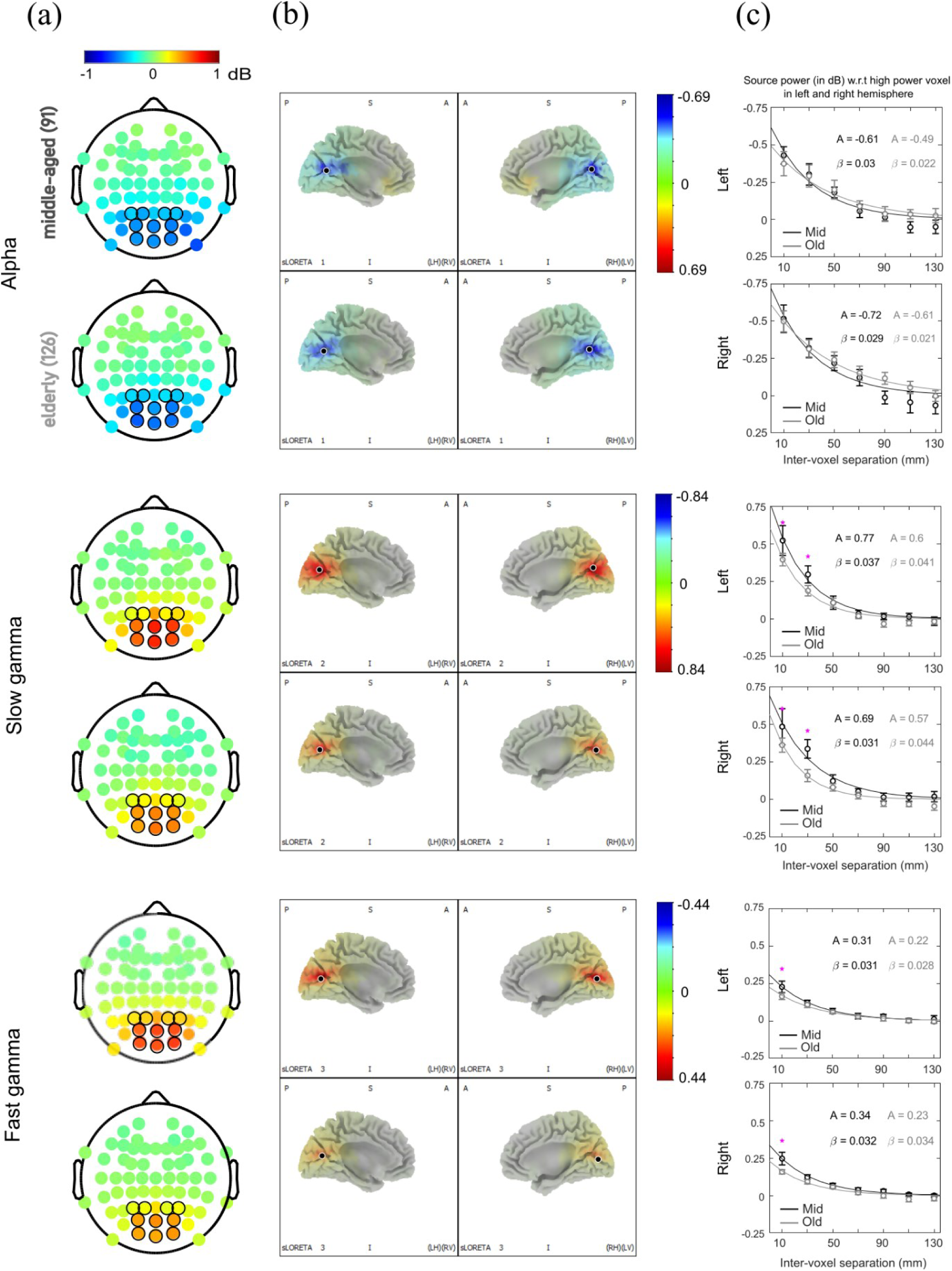
Source distribution of change in power across subjects in middle-aged and elderly age groups. (a) Median change in power scalp plots across middle-aged (age<65; N=91) and elderly (N=127) subjects, in alpha, slow gamma, and fast gamma over the rows. The labels for middle-aged and elderly are interchangeable with mid and old. (b) Source distribution of change in power across a medial sagittal cross-section in left and right hemispheres for middle-aged on top and elderly on the bottom, for the three frequency bands. The black dot represents the voxel with maximum activity, used as a reference for the power fall-off with inter-voxel separation plot in panel (c). (c) On the top is the change in source power with inter-voxel separation for middle-aged (black trace) and elderly (grey trace) groups in the left hemisphere. The data is binned over seven bins with a bin width of 20 mm centred between [0⎯140] mm, with 20 mm spacing. The median of the binned data (see Methods) across subjects, along with bootstrapped SEM, is plotted as an error bar in each of the bins, and those with significant power differences across the age groups are indicated with an asterisk. Medians are fit with an exponential decay function (see Methods). The A and β parameters are reported in the figure inset for left and right hemisphere plots. On the bottom is the similar plot, but for the voxels in the right hemisphere. For the alpha power the Y axis is reversed.

### Estimating age dependency of source extent

The source power fall-off parameters for each of the frequency ranges were checked for age dependency through two methods. The first approach was by dividing the subjects in two age groups - middle-aged (age<65; N=91) and elderly (>=65; N=121), as done in our previous studies (Murty *et al*., 2020), and comparing the median parameters across groups. This approach allowed us to view the median power distribution for each group (shown in Figure 2). We also plotted the average power across inter-voxel separation in equal interval bins along with median SEM, defined as the standard deviation of bootstrapped medians over N = 1000. This was used to display the variation of fall-off parameters across the age groups within each inter-voxel separation bin (Figure 4c). However, artificial binning of a continuous parameter such as age into arbitrary groups has several issues (Naggara *et al*., 2011). We therefore used a second approach using a linear regression model with all the subjects considered together disregarding any arbitrary age group. The fall-off parameters (A and *β*) were individually modelled to depend on the subject age as follows, in the Wilkinson notation:

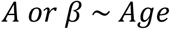

The model was fitted for the source power fall-off parameters on both left and the right hemispheres. Coefficient for age was tested for significance based on the t-statistic. To whether the changes in the Amplitude and decay parameters were simply due to the weakening of power in the sensor space as reported in the previous study (Murty *et al*., 2020), we computed the average change power (across the ten occipital electrodes listed above) and included it as another dependent variable along with the subject age in a multiple linear regression model.

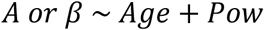

Along with the regression fits, we also plotted average power across age in equal interval bins with along with median SEM. This was used to display the variation of fall-off parameters with each age bin (Figure 5).

**Figure 5.**
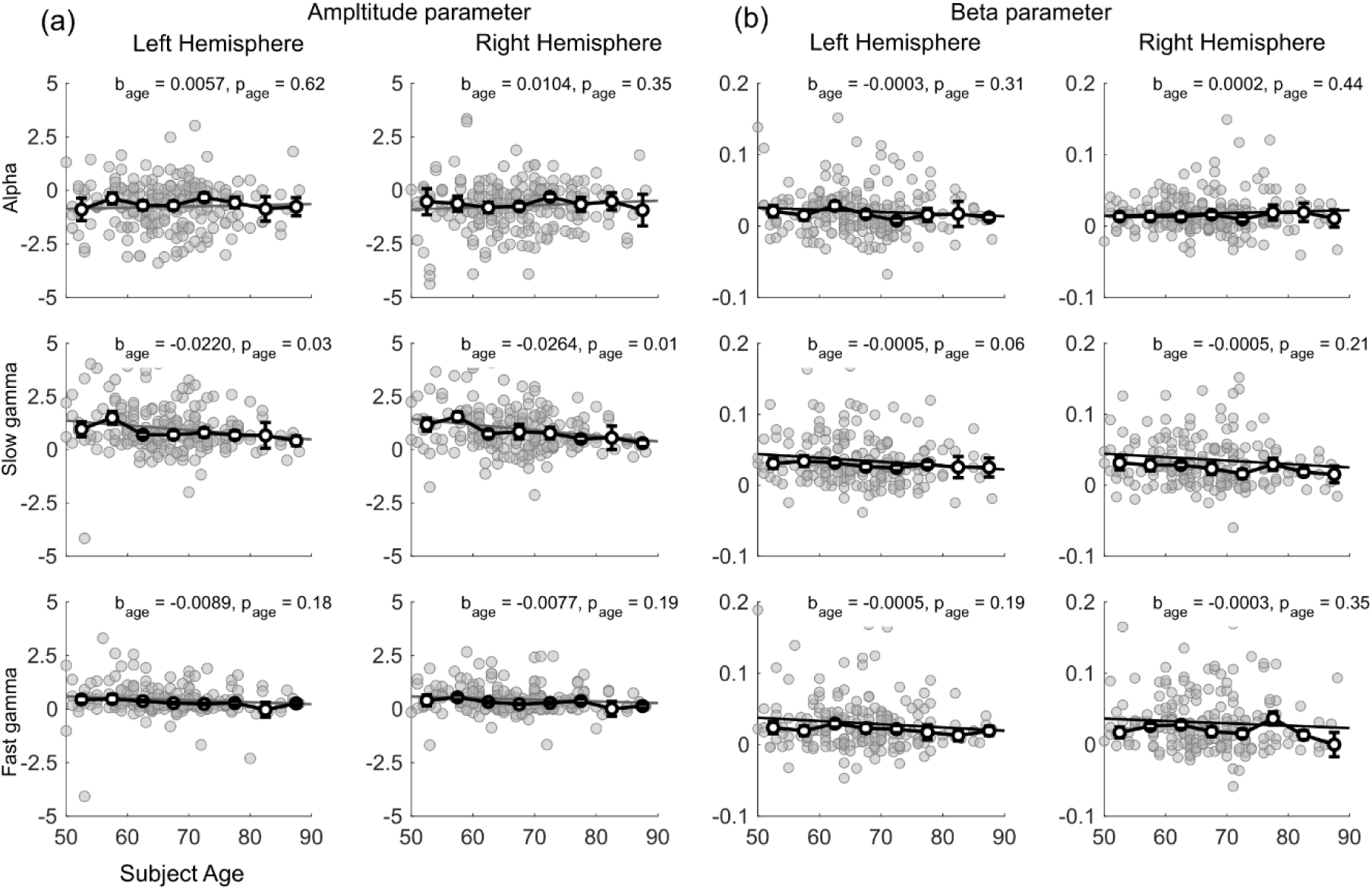
Age dependency of cortical power fall-off parameters. **(a)** The amplitude parameter estimated for each subject based on exponential decay function fitted over the cortical power is plotted against subject age, for the left and right hemispheres, within alpha, slow gamma, and fast gamma over the rows. The line plot depicts median value, within uniformly spaced bins spanning 5 years each. The error bar conveys the bootstrapped median SEM (Standard Error of Median) (refer Methods). The linear regression coefficient for age variable, with respective p-value of significance is indicated for each of the frequency bands. **(b)** The decay parameter for each subject is plotted against age similar to (a).

### Statistical Analysis

For group comparisons, non-parametric test like the Kruskal-Wallis was used to check the changes in amplitude and decay parameters. The linear regression coefficients were tested for significance based on the t-statistic.

### Data Availability Statement

Codes are freely available at https://github.com/supratimray/SourceLocalizationProject. The raw data are openly available in the OSF repository at https://osf.io/ebryn/.

## Results

First, we describe the eLORETA source localization technique for an example subject (Figure 2). Time-frequency change in power spectrogram (Figure 2a), which shows the change in power relative to a pre-stimulus baseline period (-500 to 0 ms), shows an increase power in both the slow gamma (20–35 Hz) and fast gamma (36–66 Hz) bands. Stimulus onset produced a strong transient response over, followed by a narrowband gamma response from ∼250 ms onwards. We therefore computed power between 250-750 ms (termed the “stimulus period”) for further analysis. To visualize the extent of slow gamma activity across brain regions, we plotted scalp maps (topoplots) for the slow gamma frequency range (Figure 2b). Next, we computed the sources using the eLORETA technique for both the baseline and the stimulus time periods. Figure 2c and 2d display the corresponding source distribution maps overlaid on a sagittal cross section of a template MRI image. The change in source power during the stimulus period relative to the baseline power revealed the sources activated by the stimulus (Figure 2e). To illustrate how source power decreased with distance from the voxel with maximal change in power, we plotted the mean change in source power as a function of voxel distance after binning the data using a 20 mm bin width over a range of 0 to 140 mm. We also fitted the data using an exponential decay function and estimated A and β parameters, which depict weakening and shrinkage of the sources (see Methods for details).

Since source localization is inherently an ill-posed problem, and because every source localization algorithm makes certain assumptions, whether an actual shrinkage in source distribution is adequately captured using our approach is not guaranteed. Further, for generating the source power versus intervoxel distance (Figure 2f), there is an inherent assumption of symmetric source decay about the peak power voxel along different dimensions, which may not be valid. Therefore, to validate the eLORETA technique used for source estimation and to test whether β can actually capture shrinkage of sources, we conducted a simulation using time-domain data from the same subject during the stimulus period (see Figure 1). In the simulation, we restricted the source model to different spatial extents— specifically 100 mm, 80 mm, 60 mm, and 40 mm (Figure 3a)—by setting the dipole magnitudes of all voxels beyond a certain distance from the seed voxel to zero, effectively “shrinking” the source space. For each spatial configuration, we reconstructed the time-domain sensor data from the source maps using forward modelling in FieldTrip, utilizing the lead field matrix computed for the recording electrode locations. We then computed the power at each electrode for the slow gamma frequency range and generated scalp maps that localized the power to the occipital region (Figure 3b). To cross-validate the source extent, we performed source localization on the reconstructed sensor data and compared the results to the original source distribution (Figure 3c). Finally, for both the original (Figure 3a) and reconstructed source distribution maps, we fitted the data as described in the Methods section to estimate the A and β parameters corresponding to the source maps (Figure 3d). We found that the source maps could adequately capture the shrinkage, leading to an increase in the β parameter as well.

Figure 4a shows the median change in power across all the electrodes as a scalp map for the two age groups, namely middle-aged and elderly in alpha, slow gamma, and fast gamma bands. Across the age groups, alpha remained unchanged, while the gamma power weakened within the occipital electrodes, as reported in our previous study (Murty *et al*., 2020). Corresponding change in source power maps are shown in Figure 4b (refer to the “Source Localization” section in Methods for details on the computation of source power). The source power map over the cortex sampled at the medial sagittal section is shown, such that it features the primary visual cortex across the left and right hemispheres. Similar to the results shown in sensor space (Figure 4a), source power in slow and fast gamma bands reduced with age, while no substantial change was observed in alpha.

To compare between the different hypotheses as indicated in Figure 1, we computed a one-dimensional plot of source power as a function of inter-voxel separation from the highest power voxel in the visual cortex, marked as a black circular patch on the source map in each hemisphere (Figure 4b). This power fall-off plot (Figure 4c) was fitted with an exponential with an Amplitude and a Decay parameter (*A* and *β*; refer “Parametrizing source distribution with exponential decay function” of Methods). We also compared the average power across bins of varying inter-voxel separation between the two age groups. While alpha suppression in source space remained unaltered with age, with no significant inter-voxel separation bin (top plot in Figure 4c), slow and fast gamma source power reduced in elderly over the middle-aged, with significant difference in inter-voxel separation bins that were close to the source bin (as indicated by asterisks in middle and bottom plots in Figure 4c).

In the source maps shown in Figure 4b, the gamma sources appeared to weaken (Figure 1a) rather than shrink (Figure 1b) across the two age groups. This was confirmed by comparing the amplitude and decay parameters across the two age groups. Amplitude parameter for gamma significantly reduced in old over the mid group: slow gamma, right hemisphere: (1.22 ± 0.14 for mid versus 0.84 ± 0.09 for old (median ± SE); *χ*^2^(1) = 5.09, n=91/126, p = 0.024; KW test); left hemisphere: 1.16 ± 0.14 for mid versus 0.86 ± 0.09 for old; *χ*^2^(1) = 3.46, n=91/126, p = 0.063; KW test). Fast gamma, right 0.55 ± 0.07 for mid versus 0.39 ± 0.06 for old; *χ*^2^(1) = 5.92, n=91/126, p = 0.015; KW test); left hemisphere: (0.53 ± 0.09 for mid versus 0.37 ± 0.06 for old; *χ*^2^(1) = 5.13, n=91/126, p = 0.024; KW test). On the other hand, decay parameters were not significantly different: slow gamma, right hemisphere: (0.038 ± 0.004 for mid versus 0.034 ± 0.004 for old (median ± SE); *χ*^2^(1) = 3.14, n=91/126, p = 0.076; KW test); left hemisphere: 0.04 ± 0.0036 for mid versus 0.03 ± 0.0032 for old; *χ*^2^(1) = 3.86, n=91/126, p = 0.049; KW test). Fast gamma, right hemisphere: 0.034 ± 0.0044 for mid versus 0.028 ± 0.0038 for old; *χ*^2^(1) = 1.49, n=91/126, p = 0.22; KW test and left hemisphere: (0.032 ± 0.004 for mid versus 0.029 ± 0.003 for old; *χ*^2^(1) = 0.38, n=91/126, p = 0.53; KW test) and). For alpha, neither amplitude nor decay parameters were significantly different.

### Age dependency of source gamma oscillations

Because group-wise analysis where a continuous variable such as age is artificially binned into arbitrary groups suffers from several issues (Naggara *et al*., 2011; Kuss, 2013), we tested the age dependency of the Amplitude and Decay parameters using linear regression modelling with each subject considered individually (refer Methods). Figure 5a depicts the amplitude parameters for both left and right hemisphere fall-off profiles. The regression coefficient for age, along with its significance is mentioned along with the scatter in each plot. The p-value relates to the t-statistic of the two-sided hypothesis test (*H*: *b* = 0), given the other terms (here the constant intercept) in the model. Consistent with groupwise analysis, the coefficient for the amplitude parameter for alpha was not significant but was significantly negative for slow gamma. For fast gamma, the coefficient was negative but did not reach significance, potentially due to the small absolute value and large variance. In contrast, the decay parameters (Figure 5b) did not change significantly for any frequency band.

### Source gamma oscillations weakening with aging is driven by sensor power

In a previous study, we found that the reduction in FC with age persisted even after accounting for variation in sensor power (Kumar et al., 2023). We therefore tested whether weakening (or potential shrinkage) of gamma sources could be observed after accounting for the variation in sensor power with age. This also allowed us to ascertain if the spatial filters applied through source reconstruction procedure contributed any additional information not explained by the sensor domain power. We accounted for the sensor power differences by including it as one of the dependent variables in a multiple linear regression model (refer Methods). We found that once sensor power was accounted for, the age dependency of slow gamma fall-off profile amplitude parameters (coefficient for age) was no longer significant (Left: *b_age_* = -0.0139, p-value = 0.14; Right: *b_age_* = -0.0162, p-value = 0.09). Overall, these results show that the gamma source power weakening is, unsurprisingly, driven by the weakening in the sensor space.

## Discussion

### Summary

We studied changes in the source distribution of visual stimulus-induced narrow-band gamma oscillations with healthy aging to better understand the reasons behind the reduction of gamma power in the sensor space as shown earlier (Murty et al., 2020). We tested different hypotheses (as shown in Figure 1) and found that slow gamma sources weaken but not shrink with age (consistent with Figure 1a), whereas alpha sources remained unchanged with age. These results were confirmed using both group-wise analysis and regression analysis.

### Previous studies on changes in source gamma power with age

A recent MEG study (Hoshi & Shigihara, 2020) on humans aging 22‒75 years reported similar changes in gamma power during Eyes Closed (EC) and Eyes Open (EO) conditions. Alpha (8- 13 Hz), beta (13-25 Hz) and high gamma (41-80 Hz) power in the source space reduced with age in the occipital areas of the brain for the EC condition, although the opposite trend was observed in rostral regions. During EO condition, only alpha and beta power (overlapping with the slow gamma band) were found to weaken with age. Similarly, changes in the EEG source space in various frequency bands with aging using eLORETA has been reported by Aoki and colleagues (2022). They reported increase in EEG spontaneous gamma source activity (during EC paradigm) with aging, in the frontoparietal and left temporal areas through correlation analysis. They suggested that the enhanced activity in the above mentioned areas may be due to the compensatory response of the cholinergic brainstem pathway, following the degradation in basal forebrain pathway with aging (Babiloni *et al*., 2006). The study reported no significant correlation of activity in gamma band within the occipital cortex with age. An EEG study on healthy aging (Perinelli *et al*., 2022) showed increase in gamma activity at the frontoparietal and left temporal areas. However, most of the studies in the literature on gamma are confined to the resting state condition, which is considered to have various shortcomings attributed to experimental settings, mental state of the subject, etc. (van Diessen *et al*., 2015). In contrast, stimulus-induced gamma oscillations are induced using certain visual stimuli such as bars and gratings in the visual cortex that have interesting characteristics such as separability from background oscillations, and characterization based on stimulus parameters. Therefore, the differences in results from previous studies could be due to the nature of gamma oscillations, since the stimulus induced gamma oscillations differ from the spontaneous gamma oscillations (Tallon-Baudry & Bertrand, 1999). In particular, the resting state power spectral density is known to “tilt” with age (Voytek *et al*., 2015; Aggarwal & Ray, 2023), leading to more power at higher frequencies due to flattening of the spectrum. However, this is a broadband response which occurs over a large frequency range and is distinct from the narrow-band gamma oscillations studied here.

### Relationship between source distribution and sensor power

Changes in the sensor space with aging may not necessarily translate to similar changes in the source space. For example, one study suggested a transformation on the source space based on the spherical harmonics basis set to improve source reconstruction, but for estimating dipole source activity, the leadfield matrix also had to be transformed into the new space defined by the spherical harmonics basis (Petrov, 2012). In other words, any general modification in source space activity, does not necessarily get reflected as a similar transformation on sensor space unless the leadfield matrix is correspondingly adjusted based on the data covariance. Another study reported that the leadfield matrix needs to be modified based on the transformation applied on the sensor data to estimate the underlying sources (Hipp & Siegel, 2015). Overall, these emphasize that the gamma weakening on sensor space may not lead to similar weakening of the gamma sources. Therefore, it is not straightforward to expect weakening in source space given that gamma weakens in sensor space.

Although the same sensor power distribution could be either due to a single localized source or multiple distributed sources, the associated FC patterns could be different. The localised source may lead to high FC between the sensors since all sensors essentially reflect the activity of a single source, whereas in the distributed source case, FC may critically depend on the extent of the coherent sources and may reduce if the source distribution shrinks. In other words, an overall weakening of gamma sources may have much less effect on FC compared to shrinkage. As shown previously, gamma FC reduced with age even when sensor power was accounted for (Kumar & Ray, 2023), which is more consistent with shrinkage rather than weakening of gamma sources. On the other hand, the relationship between power and FC may depend on factors beyond just source distribution. Indeed, FC parameters may depend more on the level of synchronization between sources as opposed to just the source extent. There are a variety of compensatory mechanisms that may play a role in synchronization. For example, Pathak and colleagues recently showed that increased inter-areal coupling can offset the age-related increase in latency (Pathak et al, 2022). Therefore, if the distributed sources synchronise more due to increased coupling, the FC will not reduce even if the source extent shrinks. Furthermore, weaker FC in distributed sources case could be due to the relative configuration of sources, in terms of their location and orientation. Power and FC are by definition incomparable and thereby, FC we cannot be solely predicted from the sensor power.

Our results instead present a simpler alternative regarding changes in stimulus-induced gamma oscillations with age. These oscillations are known to arise due to the interaction of excitatory and inhibitory interneurons, which forms a close network (Bartos *et al*., 2007). With aging, only the efficacy of this network reduces, leading to weaker oscillations, but not the spatial extent of the network itself. Reduction in FC could be due to a reduction in synchronization within the network. Although our source analysis provides more information about potential changes in the network, more advanced methods for studying functional connectivity and synchronization as well as detailed network models of gamma oscillations are needed to understand how these networks change with healthy aging and the effect of these changes on brain function.

### Limitations of the study

One limitation of this study is that the subjects were asked to simply fixate during the experiment, and hence it is not possible to directly gauge vigilance or level of engagement in this task paradigm. There are several reasons for not employing an attention task, such as the one employed recently (Das *et al*., 2024) to control/measure the level of vigilance. First, these tasks are complicated and usually of long duration and likely to be extremely tiring for elderly individuals. Second, the stimulus-induced gamma oscillations studied here are induced strongly using full-screen gratings but are generally much weaker for non-foveal stimuli are commonly used in spatial attention tasks (see Figure 3 of Das *et al*., 2024). Third, since both alpha and gamma power depend on the attentional/vigilance level that is likely to wane with age, imposing a task structure that has an explicit attention component may lead to more drastic changes in alpha/gamma power due to the differences in cognitive abilities with age instead of changes in brain circuitry with ageing.

Could the weakening of gamma sources with age as reported here be simply due to differences in attention or other cognitive differences? Although it is not possible to rule this factor out completely, we think it is unlikely for two reasons. First, the fixation task involved eye tracking, and several measures related to eye movements, such as eye positions, micro-saccade rates, and variation in pupil diameter etc have been studied earlier and were found to be comparable across groups (see Figure 7 of Murty *et al*., 2020). Second, and more importantly, in EEG, attention has a much stronger effect on alpha than gamma power (Das *et al*., 2024), but alpha suppression was comparable across the two age groups.

Another limitation of this study is that the source space analysis was conducted for the EEG data using a template head model. Due to constraints in the subject recording protocol, individual MRI data (T1-weighted structural scans) were not available, preventing the use of subject-specific head models for more accurate source reconstructions. Since source localization relies on regularization techniques to solve the ill-posed inverse problem, the absence of subject-specific anatomical information—such as that provided by a T1 scan— limits the accuracy of deeper brain region estimates. Several studies have highlighted the relationship between healthy aging and brain structural properties, including cortical thickness, bone density, and scalp impedance (Chen *et al*., 2010; Wendel, 2010; McCann *et al*., 2019; McCann & Beltrachini, 2021). However, due to the lack of individual MRI data, we were unable to examine whether gamma source weakening was influenced by these structural brain parameters.

## Funding disclosure and Acknowledgements

This work was supported by Tata Trusts Grant, Wellcome Trust/DBT India Alliance (Senior fellowship IA/S/18/2/504003 to SR), and DBT-IISc Partnership Programme.

## Abbreviations

AD: Alzheimer’s Disease
EEG: Electroencephalogram
MEG: Magneto-encephalogram
fMRI: Functional Magnetic Resonance Imaging
FC: Functional Connectivity
PBL: Power spectra during Baseline duration
PST: Power spectra during Stimulus duration
PSD: Power spectral density
SG: Slow Gamma
FG: Fast Gamma
eLORETA: exact Low-resolution Tomography Analysis
K-W: Kruskal-Wallis test
Median SEM: Standard of Error Median

## References

Aggarwal, S. & Ray, S. (2023) Slope of the power spectral density flattens at low frequencies (<150 Hz) with healthy aging but also steepens at higher frequency (>200 Hz) in human electroencephalogram. Cereb. Cortex Commun., 4, tgad011.

Anders, M.F. & Kristine, B.W. (2010) Structural Brain Changes in Aging: Courses, Causes and Cognitive Consequences. Rev. Neurosci., 21, 187–222.

Aoki, Y., Hata, M., Iwase, M., Ishii, R., Pascual-Marqui, R.D., Yanagisawa, T., Kishima, H., & Ikeda, M. (2022) Cortical electrical activity changes in healthy aging using EEG- eLORETA analysis. Neuroimage Rep., 2, 100143.

Aoki, Y., Takahashi, R., Suzuki, Y., Pascual-Marqui, R.D., Kito, Y., Hikida, S., Maruyama, K., Hata, M., Ishii, R., Iwase, M., Mori, E., & Ikeda, M. (2023) EEG resting-state networks in Alzheimer’s disease associated with clinical symptoms. Sci. Rep., 13, 1– 10.

Babiloni, C., Binetti, G., Cassarino, A., Dal Forno, G., Del Percio, C., Ferreri, F., Ferri, R., Frisoni, G., Galderisi, S., Hirata, K., Lanuzza, B., Miniussi, C., Mucci, A., Nobili, F., Rodriguez, G., Luca Romani, G., & Rossini, P.M. (2006) Sources of cortical rhythms in adults during physiological aging: A multicentric EEG study. Hum. Brain Mapp., 27, 162–172.

Babiloni, C., Carducci, F., Lizio, R., Vecchio, F., Baglieri, A., Bernardini, S., Cavedo, E., Bozzao, A., Buttinelli, C., Esposito, F., Giubilei, F., Guizzaro, A., Marino, S., Montella, P., Quattrocchi, C.C., Redolfi, A., Soricelli, A., Tedeschi, G., Ferri, R., Rossi-Fedele, G., Ursini, F., Scrascia, F., Vernieri, F., Pedersen, T.J., Hardemark, H.G., Rossini, P.M., & Frisoni, G.B. (2013) Resting state cortical electroencephalographic rhythms are related to gray matter volume in subjects with mild cognitive impairment and Alzheimer’s disease. Hum. Brain Mapp., 34.

Bartos, M., Vida, I., & Jonas, P. (2007) Synaptic mechanisms of synchronized gamma oscillations in inhibitory interneuron networks. Nat. Rev. Neurosci., 8, 45–56.

Borghini, G., Candini, M., Filannino, C., Hussain, M., Walsh, V., Romei, V., Zokaei, N., & Cappelletti, M. (2018) Alpha Oscillations Are Causally Linked to Inhibitory Abilities in Ageing. J. Neurosci., 38, 4418–4429.

Chen, F., Hallez, H., & Staelens, S. (2010) Influence of skull conductivity perturbations on EEG dipole source analysis. Med. Phys.,.

Das, A., Nandi, N., & Ray, S. (2024) Alpha and SSVEP power outperform gamma power in capturing attentional modulation in human EEG. Cereb. Cortex, 34, bhad412.

Delorme, A. & Makeig, S. (2004) EEGLAB: an open source toolbox for analysis of single-trial EEG dynamics including independent component analysis. J. Neurosci. Methods, 134, 9–21.

Dennis, E.L. & Thompson, P.M. (2014) Functional Brain Connectivity Using fMRI in Aging and Alzheimer’s Disease. Neuropsychol. Rev., 24, 49–62.

Gianotti, L.R.R., Künig, G., Faber, P.L., Lehmann, D., Pascual-Marqui, R.D., Kochi, K., & Schreiter-Gasser, U. (2008) Rivastigmine effects on EEG spectra and three-dimensional LORETA functional imaging in Alzheimer’s disease. Psychopharmacology (Berl.), 198, 323–332.

Halder, T., Talwar, S., Jaiswal, A.K., & Banerjee, A. (2019) Quantitative Evaluation in Estimating Sources Underlying Brain Oscillations Using Current Source Density Methods and Beamformer Approaches. eNeuro, 6.

Hara, Y., Rapp, P.R., & Morrison, J.H. (2012) Neuronal and morphological bases of cognitive decline in aged rhesus monkeys. Age,.

Hassler, U., Trujillo Barreto, N., & Gruber, T. (2011) Induced gamma band responses in human EEG after the control of miniature saccadic artifacts. NeuroImage, 57, 1411–1421.

Hipp, J.F. & Siegel, M. (2015) Accounting for Linear Transformations of EEG and MEG Data in Source Analysis. PLOS ONE, 10, e0121048.

Hoshi, H. & Shigihara, Y. (2020) Age- and gender-specific characteristics of the resting-state brain activity: a magnetoencephalography study. Aging, 12, 21613–21637.

Jatoi, M.A., Kamel, N., Malik, A.S., Faye, I., & Begum, T. (2014) A survey of methods used for source localization using EEG signals. Biomed. Signal Process. Control, 11, 42–52.

Kaboodvand, N., Karimi, H., & Iravani, B. (2024) Preparatory activity of anterior insula predicts conflict errors: integrating convolutional neural networks and neural mass models. Sci. Rep., 14, 16682.

Knyazeva, M.G. (2021) Chapter 31 - Alpha rhythms: what they are and how they alter with aging. In Martin, C.R., Preedy, V.R., & Rajendram, R. (eds), Factors Affecting Neurological Aging. Academic Press, pp. 349–359.

Kumar, W.S., Manikandan, K., Murty, D.V.P.S., Ramesh, R.G., Purokayastha, S., Javali, M., Rao, N.P., & Ray, S. (2022) Stimulus-Induced Narrowband Gamma Oscillations are Test–Retest Reliable in Human EEG. Cereb. Cortex Commun., 3, tgab066.

Kumar, W.S. & Ray, S. (2023) Healthy ageing and cognitive impairment alter EEG functional connectivity in distinct frequency bands. Eur. J. Neurosci., 58, 3432–3449.

Kuss, O. (2013) The danger of dichotomizing continuous variables: A visualization. Teach. Stat., 35, 78–79.

Lehmann, D., Faber, P.L., Tei, S., Pascual-Marqui, R.D., Milz, P., & Kochi, K. (2012) Reduced functional connectivity between cortical sources in five meditation traditions detected with lagged coherence using EEG tomography. Neuroimage, 60, 1574–1586.

Lopes da Silva, F. (2013) EEG and MEG: Relevance to Neuroscience. Neuron, 80, 1112–1128.

Mandal, P.K., Banerjee, A., Tripathi, M., & Sharma, A. (2018) A Comprehensive Review of Magnetoencephalography (MEG) Studies for Brain Functionality in Healthy Aging and Alzheimer’s Disease (AD). Front. Comput. Neurosci., 12.

Mazziotta, J., Toga, A., Evans, A., Fox, P., Lancaster, J., Zilles, K., Woods, R., Paus, T., Simpson, G., Pike, B., Holmes, C., Collins, L., Thompson, P., MacDonald, D., Iacoboni, M., Schormann, T., Amunts, K., Palomero-Gallagher, N., Geyer, S., Parsons, L., Narr, K., Kabani, N., Le Goualher, G., Boomsma, D., Cannon, T., Kawashima, R., & Mazoyer, B. (2001) A probabilistic atlas and reference system for the human brain: International Consortium for Brain Mapping (ICBM). Philos. Trans. R. Soc. B Biol. Sci.,.

McCann, H. & Beltrachini, L. (2021) Impact of skull sutures, spongiform bone distribution, and aging skull conductivities on the EEG forward and inverse problems. J. Neural Eng.,.

McCann, H., Pisano, G., & Beltrachini, L. (2019) Variation in Reported Human Head Tissue Electrical Conductivity Values. Brain Topogr.,.

Murty, D.V., Manikandan, K., Kumar, W.S., Ramesh, R.G., Purokayastha, S., Nagendra, B., M. L., A., Balakrishnan, A., Javali, M., Rao, N.P., & Ray, S. (2021) Stimulus-induced gamma rhythms are weaker in human elderly with Mild Cognitive Impairment and Alzheimer’s Disease. eLife, 10, e61666.

Murty, D.V.P.S., Manikandan, K., Kumar, W.S., Ramesh, R.G., Purokayastha, S., Javali, M., Rao, N.P., & Ray, S. (2020) Gamma oscillations weaken with age in healthy elderly in human EEG. NeuroImage, 215, 116826.

Murty, D.V.P.S. & Ray, S. (2022) Stimulus-induced Robust Narrow-band Gamma Oscillations in Human EEG Using Cartesian Gratings. Bio-Protoc., 12, e4379–e4379.

Murty, D.V.P.S., Shirhatti, V., Ravishankar, P., & Ray, S. (2018) Large Visual Stimuli Induce Two Distinct Gamma Oscillations in Primate Visual Cortex. J. Neurosci., 38, 2730– 2744.

Naggara, O., Raymond, J., Guilbert, F., Roy, D., Weill, A., & Altman, D.G. (2011) Analysis by Categorizing or Dichotomizing Continuous Variables Is Inadvisable: An Example from the Natural History of Unruptured Aneurysms. *Am*. J. Neuroradiol., 32, 437–440.

Oostenveld, R., Fries, P., Maris, E., & Schoffelen, J.-M. (2011) FieldTrip: Open Source Software for Advanced Analysis of MEG, EEG, and Invasive Electrophysiological Data. Comput. Intell. Neurosci., 2011, 156869.

Pagano, S., Fait, E., Monti, A., Brignani, D., & Mazza, V. (2015) Electrophysiological Correlates of Subitizing in Healthy Aging. PLOS ONE, 10, e0131063.

Pascual-Marqui, R.D. (2007) Discrete, 3D distributed, linear imaging methods of electric neuronal activity. Part 1: exact, zero error localization. ArXiv Prepr. ArXiv07103341,.

Pascual-Marqui, R.D. (2009) Theory of the EEG inverse problem. *No Title*, 121.

Pascual-Marqui, R.D., Lehmann, D., Koukkou, M., Kochi, K., Anderer, P., Saletu, B., Tanaka, H., Hirata, K., John, E.R., & Prichep, L. (2011) Assessing interactions in the brain with exact low-resolution electromagnetic tomography. Philos. Trans. R. Soc. Math. Phys. Eng. Sci., 369, 3768–3784.

Pathak, A., Sharma, V., Roy, D., & Banerjee, A. (2022) Biophysical mechanism underlying compensatory preservation of neural synchrony over the adult lifespan. *Commun*. Biol., 5, 1–12.

Pazo-Álvarez, P., Amenedo, E., Lorenzo-López, L., & Cadaveira, F. (2004) Effects of stimulus location on automatic detection of changes in motion direction in the human brain. Neurosci. Lett., 371.

Perinelli, A., Assecondi, S., Tagliabue, C.F., & Mazza, V. (2022) Power shift and connectivity changes in healthy aging during resting-state EEG. NeuroImage, 256, 119247.

Petrov, Y. (2012) Harmony: EEG/MEG Linear Inverse Source Reconstruction in the Anatomical Basis of Spherical Harmonics. PLOS ONE, 7, e44439.

Porcaro, C., Balsters, J.H., Mantini, D., Robertson, I.H., & Wenderoth, N. (2019) P3b amplitude as a signature of cognitive decline in the older population: An EEG study enhanced by Functional Source Separation. NeuroImage, 184, 535–546.

Riddle, D.R. (2007) Brain Aging: Models, Methods, and Mechanisms. CRC Press.

Rossini, P.M., Rossi, S., Babiloni, C., & Polich, J. (2007) Clinical neurophysiology of aging brain: From normal aging to neurodegeneration. Prog. Neurobiol., 83, 375–400.

Scally, B., Burke, M.R., Bunce, D., & Delvenne, J.-F. (2018) Resting-state EEG power and connectivity are associated with alpha peak frequency slowing in healthy aging. Neurobiol. Aging, 71, 149–155.

Tallon-Baudry, C. & Bertrand, O. (1999) Oscillatory gamma activity in humans and its role in object representation. Trends Cogn. Sci., 3, 151–162.

van Diessen, E., Numan, T., van Dellen, E., van der Kooi, A.W., Boersma, M., Hofman, D., van Lutterveld, R., van Dijk, B.W., van Straaten, E.C.W., Hillebrand, A., & Stam, C.J. (2015) Opportunities and methodological challenges in EEG and MEG resting state functional brain network research. Clin. Neurophysiol., 126, 1468–1481.

Voytek, B., Kramer, M.A., Case, J., Lepage, K.Q., Tempesta, Z.R., Knight, R.T., & Gazzaley, A. (2015) Age-Related Changes in 1/f Neural Electrophysiological Noise. J. Neurosci., 35, 13257–13265.

Wendel, K. (2010) The Influence of Tissue Conductivity and Head Geometry on EEG Measurement Sensitivity Distributions.

